# Lesion aware automated processing pipeline for multimodal neuroimaging stroke data and The Virtual Brain (TVB)

**DOI:** 10.1101/2023.08.28.555078

**Authors:** Patrik Bey, Kiret Dhindsa, Amrit Kashyap, Michael Schirner, Jan Feldheim, Marlene Bönstrup, Robert Schulz, Bastian Cheng, Götz Thomalla, Christian Gerloff, Petra Ritter

## Abstract

**Background:** Processing stroke magnetic resonance imaging (MRI) brain data can be susceptible to lesion-based abnormalities. In this study we developed and validated the Lesion Aware automated Processing Pipeline (LeAPP) that incorporates mitigation measures, improving volumetric and connectomics outputs compared to current standards in automated MRI processing pipelines.

**Methods:** Building upon the Human Connectome Project (HCP) minimal processing pipeline, we introduced correction measures, such as cost-function masking and virtual brain transplant, and extended functional and diffusion processing to match acquisition protocols often found in a clinical context. A total of 51 participants (36 stroke patients (65.7±12.96 years, 18 female) and 15 healthy controls (69.2±7.4 years, 7 female)) were processed across four time points for patients (3-5, 30-40, 85-95, 340-380 days after stroke onset) and one time point for controls. Artificially lesioned brains (N=82), derived from healthy brains and informed by real stroke lesions were created, thus generating ground-truth data for validation. The processing pipeline and validation framework are available as containerized open-source software. Reconstruction quality has been quantified on whole brain level and for lesion affected and unaffected regions-of-interest (ROIs) using metrics like dice score, volume difference and center-of-gravity distance. Global and local level connectome reconstruction was assessed using node strength, node centrality and clustering coefficient.

**Results:** The new pipeline LeAPP provides close reconstructions of the ground truth. Deviations in reconstructed averaged whole brain node strength and all ROI based volume and connectome metrics were significantly reduced compared to the HCP pipeline without stroke specific mitigation measures.

**Conclusions:** LeAPP improves reconstruction quality of multimodal MRI processing for brain parcellation and structural connectome estimation significantly over the non-adapted HCP in the presence of lesions and provides a robust framework for diffusion and functional image processing of clinical stroke data. This novel open-source automated processing pipeline contributes to a development towards reproducible research.

**Graphical abstract:** 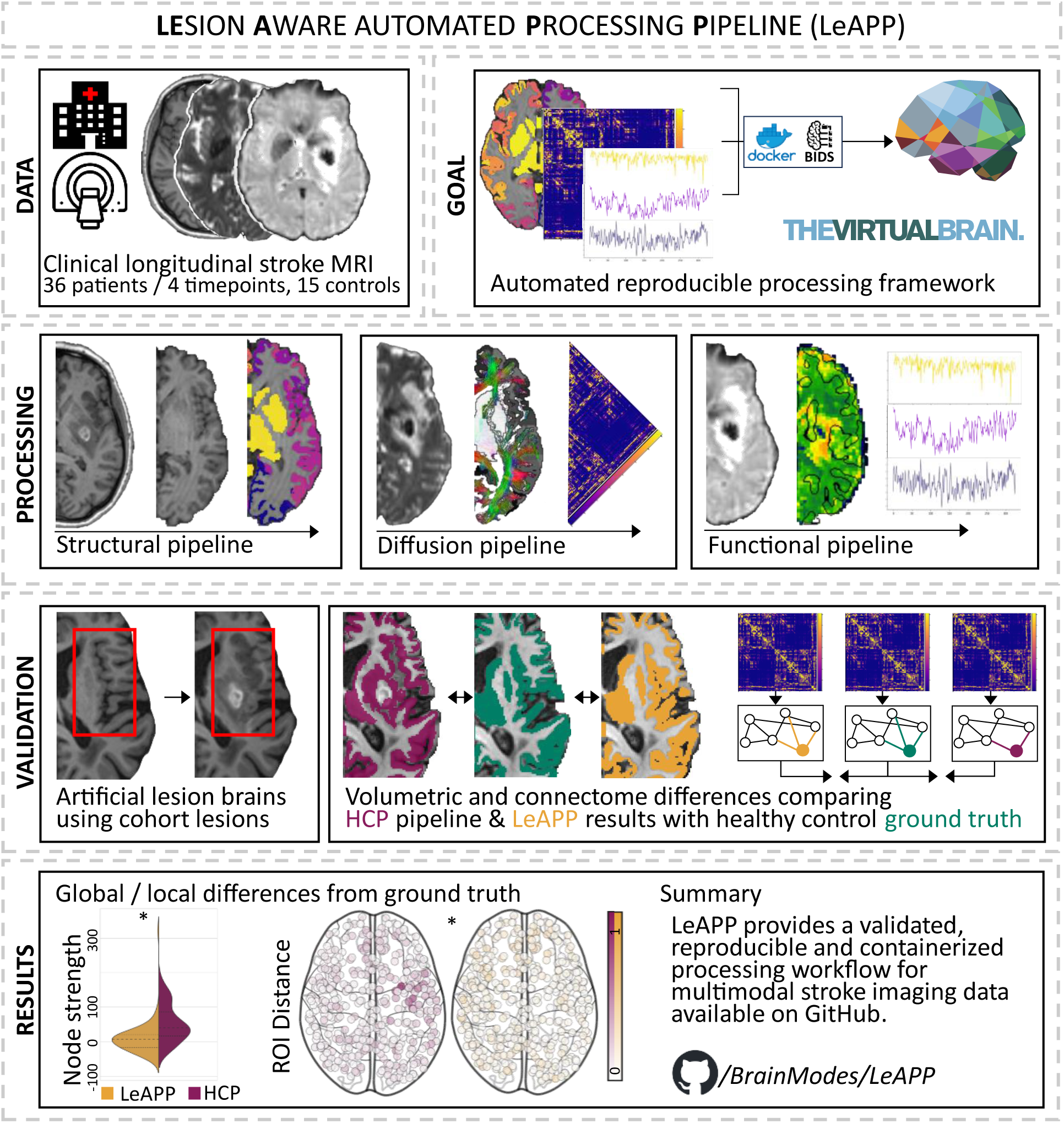

## Introduction

Processing brain imaging data from stroke patients presents challenges for existing workflows used for standardized processing of multimodal brain imaging data^1^. Particularly in magnetic resonance imaging (MRI), large lesions can cause distortions and result in performance loss and failure of common processing steps, such as surface reconstruction, brain tissue segmentation and co-registration of different images^1,2^. Established neuroimage processing pipelines often lead to failure or low-quality outputs when applied to such data sets^3^. Most stroke related MRI studies therefore rely on customized tools and individually crafted solutions^4^, report insufficient details on lesion specific adjustments made^2^ or use standard processing pipelines originally developed for healthy subjects with potentially poor processing outcomes. Therefore, a fully automated and reproducible image processing pipeline that correctly accounts for abnormalities induced by stroke lesions is needed by the scientific and clinical stroke community^3^. To address this need, we developed and validated a lesion aware automated containerized processing pipeline called LeAPP that performs structural, diffusion and functional MRI processing. We applied this novel pipeline to a longitudinal dataset of stroke patients and healthy controls^5,6^. We used the human connectome project (HCP) minimal processing pipeline^7^ as the basis for our workflow and added specific processing steps and already established correction methods to cope with the challenges of stroke brains.

Those additional methods include (1) cost function masking (CFM)^8,9^ for all coregistration steps, which restricts the fitting optimization of coregistration to healthy brain tissue, therefore minimizing lesion impact and improving overall accuracy, and (2) virtual brain transplant (VBT)^10,11^, which aims to approximate the underlying healthy tissue at the focal lesion by using contralesional hemisphere information, enabling downstream processing such as segmentation and surface extraction. Both methods are available within existing frameworks (e.g. FSL flirt variable for cost function masking or BCBToolkit^12^ for performing enantiomorphic normalization) but are not integrated in a fully automated and comprehensive processing pipeline leading to the need for manual and non-reproducible processing steps.

In addition, we performed an extensive validation of LeAPP and demonstrated significant improvement in reconstructing the underlying subject specific anatomy over the processing results obtained with the HCP pipeline. Furthermore, our pipeline automatically creates data that can serve as input to the virtual brain (TVB)^13,14^, and that can be used to construct patient specific whole brain models, thus facilitating further research into underlying mechanisms and disease patterns of stroke^15^.

## Methods

Where applicable, this study fulfills the TRIPOD checklist (Supplementary Table S4) for description of the development and validation of the pipeline. The authors declare that the source code required for replicating the present study is made available and the software is disseminated via containerized packages for the processing and validation pipelines described here (www.github.com/brainmodes/LeAPP).

### Patients

The here processed human data were acquired at University Hospital Hamburg Eppendorf ^6^. The present study was approved by the Ethics Committee of the Charité (EA1/222/22) and written consent was given by all participants for data acquisition for the original study which was approved by the ethical board of the Hamburg University Hospital (PV3777). A total of 51 participants (36 stroke patients (mean age (standard deviation) = 65.7 (12.96) years, 18 female) and 15 healthy controls (mean age (standard deviation) = 69.2 (7.4) years, 7 female)) with complete datasets (see definition in **Figure 1(D)**) at timepoint 1 (acute phase 3-5 days post stroke onset) were selected from a larger sample (n=80 (43 male). Where available, longitudinal data acquired over up to three follow up dates (30-40 days, 85-95 days and 340-380 days post stroke onset) were included as well. The data set included structural MRI, task-based functional MRI as well as diffusion weighted MRI data^5^ (full description of the included data set in supplementary **table S2**). Initial inclusion criteria for stroke patients were first ever ischemic stroke, hand-motor deficit without accompanying other functional deficits and no MRI contra indicators. Input and result (derivative) data will be made discoverable via the EBRAINS knowledge graph (https://search.kg.ebrains.eu/). Sharing of these data is subject to the EU General Data Protection Regulations (GDPR) and requires the establishment of a research purpose of processing, conclusion of EU standard contractual clauses between controllers and processors and a data protection impact assessment (DPIA) approved by the relevant institutional data protection officers.

**Figure 1:**
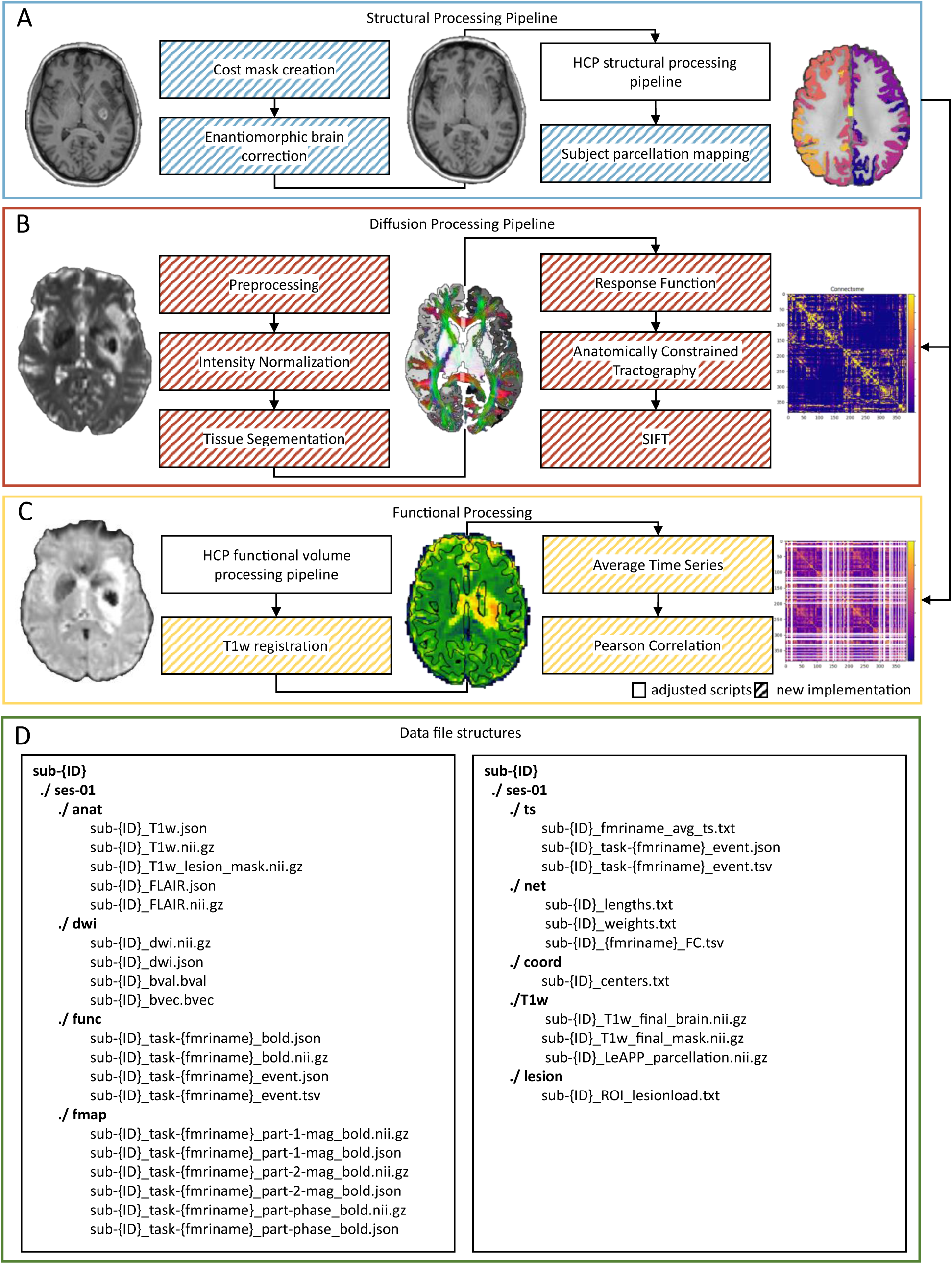
Lesion aware automated processing pipeline (LeAPP) overview. The major processing steps and adjustments for processing of structural (blue), diffusion (red) and functional MRI (orange) as implemented in LeAPP as well as required input files and TVB – ready outputs (green). Structural processing (**A**) performs the three main adjustments implemented in LeAPP build around the baseline HCP structural processing step. The output required for subsequent processing are fully processed anatomical images in native T1w and MNI space, the created brain parcellation volume and lesion properties table containing local lesion load for all affected ROIs. The diffusion processing workflow (**B**) performs the main preprocessing steps, tractography and connectome creation using the output from structural processing. (**C**) Functional preprocessing is performed based on HCP functional volume-based processing step followed by registration, average time series extraction and functional connectivity matrix creation. (**D**) Data input is following BIDS standard for all input modalities (left) including T1w and FLAIR images, DWI images with corresponding bval and bvec files, functional MRI with task event files and corresponding fieldmaps. The main results (right) are stored in a new directory including structural and functional connectomes, parcellation volumes, final T1w images and lesion mask as well as average ROI based time series with task event files following the BIDS computational modelling extension proposal and can be integrated into the virtual brain (TVB) simulation platform for individual whole brain network simulations.

### Pipeline

Image processing was implemented in four steps, structural image processing, diffusion image processing, functional image processing and output preparation. Lesion specific adjustments were integrated based on a priori defined lesion mask. Such masks have been manually drawn using ITK Snap^16^ incorporating T1w and FLAIR image information. Each processing step, as illustrated in **Figure 1**, was specifically adapted for the challenges of processing stroke lesion MRI data as follows:

*Structural image pipeline* (**Figure 1 (A)**) extends the HCP minimal processing pipeline’s structural processing^7^ to be more robust and broadly applicable when processing MRI data in the presence of stroke lesions. First, it incorporates CFM in all registration steps, which is necessary to ensure accurate registration of lesioned images^9^ by restricting the fitting of the coregistration to voxels outside of the provided lesion mask. Three additional steps are included that add to the HCP structural processing pipeline: (1) the automated creation of lesion masks, (2) the first fully automated implementation of VBT (see Supplementary Material) and (3) the creation of subject specific brain parcellations. For all structural input modalities including T1w, T2w and/or FLAIR and reference images for standardized MNI space, including MNI1mm and MNI2mm template images, undergoing realignment to other image volumes during structural processing commonly referred to as coregistration, the initially provided lesion mask is registered. Such an alignment is performed, by first fitting the linear affine coregistration using the corresponding base image (e.g., T1w image for a lesion mask in T1w space) and applying the resulting transformation to the binary mask followed by Boolean inversion. This ensures spatially aligned lesion masks. For subsequent CFM across all coregistration steps, the lesion voxels that are encoded as zeros in the aligned mask, were therefore excluded from the registration fitting process. Following ^10^ we integrated VBT by imputation of contralesional hemisphere signal into the lesion area (**Supplementary Figure S1**). This is achieved by first performing midline alignment of the input image. To this end the coregistration to the mirror image is computed and the resulting transformation matrix is halved creating a transformation that moves each point in alignment with the midline, as it represents half the distance to the mirrored point in the reference image.

The initial lesion mask for the corresponding modality is also aligned at the midline and inverted. To extract the healthy brain signal from the mirror image and remove the lesion region, the aligned lesion mask is multiplied with the midline aligned mirror image and the inverse to the aligned input image. Before extraction of both lesion and healthy signal, the border of the corresponding lesion mask image is smoothed by applying a Gaussian kernel (with 2mm full width at half maximum) to ensure a more seamless integration into neighboring voxels during imputation. The healthy signal is then added to the lesion free input image to create an approximation of the underlying healthy anatomy. To complete the implementation of VBT, the inverse midline transform is applied to create the transplanted image in native patient space of the original input image. This is applied to both T1w and corresponding T2 or FLAIR images. Following VBT, the CFM adjusted structural processing steps PreFreeSurfer, FreeSurfer and PostFreeSurfer of the HCP pipeline are initiated. Subject-specific brain parcellations are created in the final step of the *structural image pipeline*. A total of 383 distinct brain regions-of-interest (ROI) are identified using a combination of the HCP-MMP1 atlas^17^ for cortical areas and FreeSurfer’s subcortical areas^18^. To ensure improved sensitivity over standard volume space atlas mapping, this study follows^19^ performing the mapping of HCP-MMP1 regions on the cortical labels created during surface extraction in fsaverage space using a previously published mapping of HCP-MMP1 annotation labels^20^. The resulting annotation files are then mapped back into virtually transplanted volume space to create an accurate parcellation of the subjects cortical and subcortical regions at lesion free areas as well as a substantiated approximation of the underlying regions at the lesion location within high resolution native subject T1w space. In a final step the parcellation image is multiplied with the final lesion mask in T1w space to extract the lesion load per ROI, defined as the number of affected voxels divided by the total number of voxels for a given ROI.

*Diffusion image pipeline* (**Figure 1 (B)**) was implemented using the MRTrix3^21^ software package. The main processing steps for the DWI pipeline are (1) preprocessing and normalization of raw input images, (2) tissue segmentation of the corresponding anatomical image, and finally (3) tractography and connectome creation. Preprocessing steps are comprised of denoising, degibbsing, eddy current and motion correction, coregistration to T1w space for distortion correction in the absence of reverse phase encoding data and bias correction. The images are subsequently intensity normalized at the group level. Each preprocessed T1w image is segmented into five tissue types using the MRtrix3 5ttgen function and the corresponding lesion mask is integrated as a pathological tissue type. This enables anatomically constrained tractography (ACT)^22^ after group level response function estimation. The default number of streamlines created is 100 million feasible tracks with the previously segmented grey matter - white matter barrier as the seed region. The individual brain parcellation created during structural processing is used to define boundaries to construct the structural connectome using the SIFT2 algorithm^23^.

*Functional image pipeline* follows the HCP’s fMRIVolume (**Figure 1 (C)**) pipeline processing step^7^ updated with CFM and performs additional steps of (1) coregistration to T1w space and (2) creation of ROI based average time series and the functional connectome. The preprocessed output volume of the adjusted fMRIVolume pipeline is linearly registered to the preprocessed subject T1w image output from the *structural pipeline* using the created single band reference image. Applying the brain parcellation image to the registered processed time series data, the ROI specific average voxel time series are extracted and correlated using Pearson correlation coefficient to create the functional connectome.

*Output* file selection creates a collection of the final processing results within a TVB ready data format (**Figure 1 (D)**). This set of files includes the processed T1w image, corresponding brain parcellation, functional connectomes and average time series for all processed tasks, structural connectome weights and tract lengths as well as a list of ROI specific lesion loads and ROI center coordinates file. The format of the output files follows the current Brain Imaging Data Structure (BIDS) computational modelling extension proposal^24^.

### Validation

Following previous studies^9,25,26^ we evaluated the performance of LeAPP by creating artificial stroke patient brains of which a ground-truth was available for statistical comparison of pipeline outputs. The ground truth consists of the processed data of the healthy controls before integration of lesion signal.

*Artificial lesion embedding* (ALE) was performed by imputation of stroke patient lesion signal into a healthy brain volume (**FIGURE 2 (A)**). The method is closely related to the above mentioned VBT method but differs in the following aspects: instead of imputation of healthy contralesional hemisphere signal, lesion signal from a different subject (stroke patient) was used to replace healthy tissue of the subject (healthy control) which in turn represented a different coregistration approach between subjects as opposed to mirror images of the same subject as in VBT.

**Figure 2:**
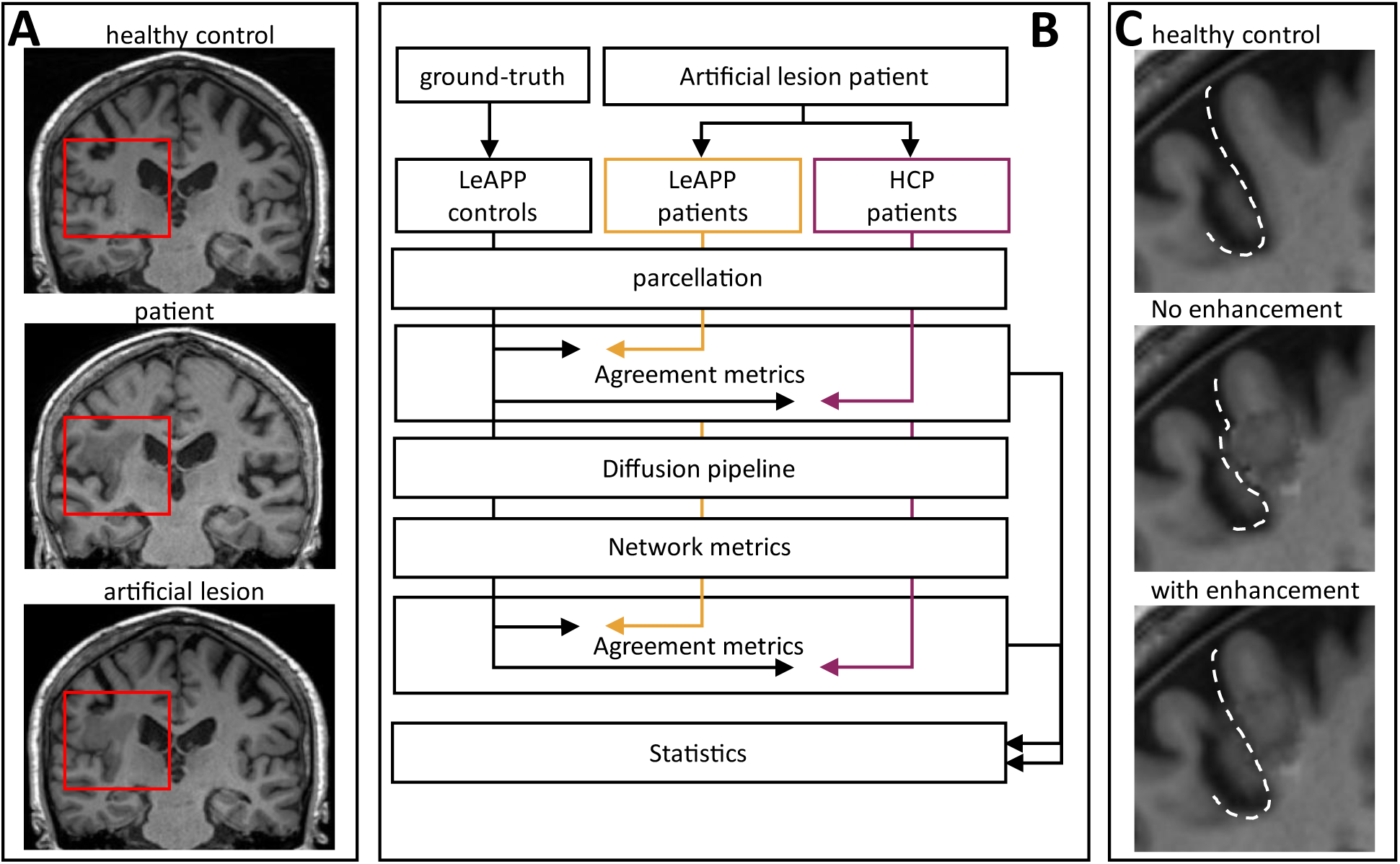
Validation framework. Example of the implemented artificial lesion embedding (ALE) pipeline (**A**) showing the original healthy control T1w image (top), the artificial lesion embedded in the same T1w(middle) and the original patient data used for extracting lesion signal (bottom). The resulting ALE data set presents a realistic approximation of a virtual stroke patient as basis for a robust validation process. (**B**) The three validation data sets (ground truth, ALE with LeAPP corrections and ALE with only baseline HCP structural processing) are first processed according to their designation followed by the LeAPP specific parcellation mapping. Based on the resulting parcellations global and local agreement measures are computed between ground truth and the ALE based results. The distributions of agreement measures are compared between LeAPP and baseline HCP based results (left). The brain parcellations are used for structural connectome creation using ground truth DWI data. A range of global and local network metrics are computed for the connectomes and agreement between metrics of ground truth and the ALE based connectomes are computed. The distributions of these agreement measures are compared between LeAPP and baseline HCP (right) (**C**) Details showing the enhancements effect during ALE of smoothing and thresholding the lesion mask to fit the underlying healthy control brain topologies more closely creating realistic artificial lesion brains.

The healthy input image was first linearly affine registered to the patient input image with CFM applied. To incorporate individual anatomical landmarks of the healthy brain in the resulting artificial stroke patient brains, such as cerebrospinal fluid of the healthy brains, the patient lesion mask was then adjusted by removing voxels whose intensity value in the registered healthy input image were below the threshold of the five percent quantile^3^, thus avoiding the imputation of lesion signal at voxels that represent for example ventricles, fissures or would span across gyri in the resulting artificial stroke patient brains. Maintaining the original landmarks of the healthy brain was necessary in order to facilitate the creation of artificial stroke patients with realistic lesion manifestations which allowed for a more thorough validation of the processing framework as opposed to unrealistic artificial stroke patients potentially introducing biases cause by unrealistic lesion properties not representative of the true patient population (**Figure 2 C**). The updated lesion mask – that excluded important landmarks - was smoothed and used to extract the lesion signal from the patient image and remove brain signal from the healthy input image. The extracted lesion signal was rescaled to fit the same scale of intensity values present in the healthy image and imputed into the adjusted healthy image, and finally, the inverse transformation of the initial coregistration to patient space was applied to create realistic artificial patient image for an existing healthy control image. This process was performed for each healthy control subject, integrating the lesion of fifteen randomly selected stroke patients resulting in 225 ALE data sets. Visual quality control was performed to ensure appropriate lesion embedding, subsequently excluding ALE combinations with lesions ranging into cerebrospinal fluid, skull, or other unfeasible areas where lesions might not manifest in that manner in stroke patient populations, representing a clear distinction as artificial. This was necessary as the random combination of subjects can lead to mismatches between source and target brain of the lesion imputation, e.g. in brain size, that cause inconsistencies in the ALE brain that could not be resolved with the previously described thresholding and smoothing. A total of 82 ALE samples remained for validation. The validation data set therefore consisted of three separate datasets (**FIGURE 2 (B)**): The original healthy control brain data used to generate the artificial patient brain data, which serves as a ground truth (LeAPP controls); the artificial stroke patient brain data corrected and processed with LeAPP (LeAPP patients); and the artificial stroke patient brain data processed using the HCP structural processing pipeline (HCP patients). We investigated the impact of the artificial stroke lesions and the corrections made by LeAPP on the final brain parcellations and network properties.

#### Reconstruction quality metrics

We next compared the performance of LeAPP and the HCP processing pipeline in recovering the ground truth brain parcellations and connectomes, which was defined as the parcellation and the connectome of the healthy brains that had been artificially distorted through the insertion of a stroke lesion. The goal, and hence quality criteria for pipeline performance, was the recovery of original parcellations and connectomes of the healthy brains – despite the artefacts introduced by the stroke lesion. We chose the following metrics to first assess the agreement of the individual brain parcellations providing information on three base errors of segmentation commonly evaluated in medical image segmentation validation: area, content and contour^27^. To this end dice coefficients, Jaccard scores, volume differences^28,29^ and Euclidean distance of the ROI center-of-gravity in voxel space were computed.

Dice coefficients are the most used metric to validate medical image segmentations and provide an estimate of the overlap of two volumes. Jaccard scores are closely correlated as another overlap-based metric of agreement^28^ approximating the base errors of area and contour. Volume differences have been computed to assess the similarity in content providing an estimate of potential distortions regarding the overall size of the compared volumes. To first validate global differences between processing modes, agreement was evaluated by comparing the binarized full brain parcellations, created during the structural processing of the ground truth controls and ALE patients, containing cortical and subcortical brain areas within a single volume mask (**Figure 3**) and computing dice coefficient, Jaccard score and volume difference. Furthermore, local differences were investigated using the non-binary individual brain parcellations, by extracting the corresponding ROI masks for both ground truth and ALE based parcellations for each ROI individually and computing dice score, volume difference and Euclidean distance between centers-of-gravity of both ROI masks.

**Figure 3.**
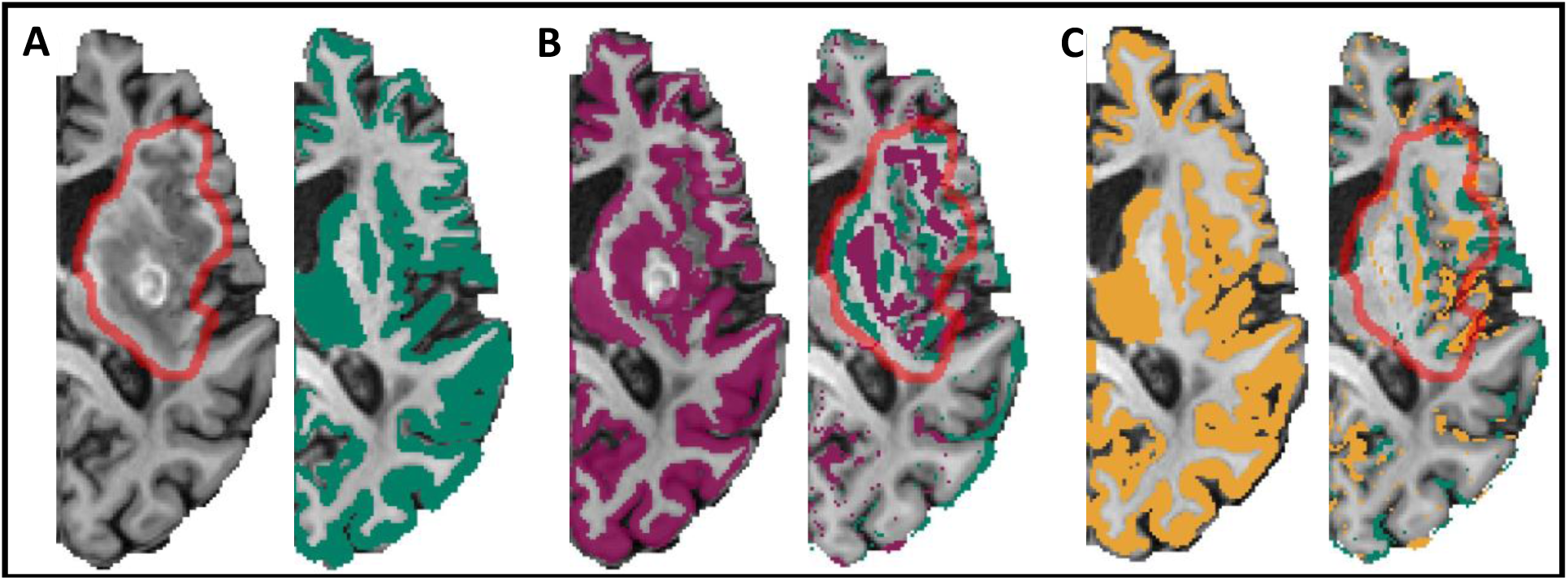
Parcellation masks. Example artificial lesion brain and corresponding Processing results. (**A**) ALE brain with imputed adjusted lesion signal (red outline) and ground truth parcellation created for healthy control used during ALE (green). (**B**) Parcellation mask created using HCP processing pipeline and parcellation mapping (purple) showing a clear reproduction of lesion topology during extraction of cortical ribbons (left). The difference between ground truth (green) and HCP (purple) with color coding voxels that are only present in the corresponding parcellation (right). (**C**) Parcellation mask created using LeAPP pipeline (yellow) and corresponding difference mask (right). While a mismatch between parcellations for LeAPP and the ground truth pertains, it shows a reduced difference at the lesion (red outline) compared to HCP.

In a second step processing mode impact on connectomes was investigated focusing on the overall connectivity and integration of the created networks. Hence node strength, centrality and clustering coefficient were computed and differences in these metrics between processing modes were defined as connectome level agreement.

Node strength provides an estimate of the overall connection of a given node, here the ROI of the brain parcellation, with all other nodes of the network, incorporating not only the existence of a connection between two nodes but the strength of the connection, here the weight of the structural connectome, as well. Betweenness centrality is widely used to assess the relative role of a node within a network in respect to efficient information exchange due to the number of shortest paths passing through the given node. Clustering coefficient provides an additional measure on the degree of the forming of local groups of nodes within the overall connectome.

In a first step again, global impact was investigated first using the averaged node strength and betweenness centrality over all ROIs as well as the clustering coefficient of the full connectome. Local differences were then evaluated by first computing the node strength and betweenness centrality for each ROI individually. Agreement was then defined as the difference in the resulting global and local measures between ground truth and ALE patient values for both processing groups (**Figure 2 B**)

### Statistical analysis

Python software package^30^ was used for statistical analysis as integrated in the containerized validation framework of this study. One-sided dependent t-tests (alpha=0.05) were computed to compare LeAPP with the HCP on both volume-based and network-based reconstruction quality metrics (**Figure 2**) reporting both the p-value for significant and t-statistic as a measure of strength in difference between the compared distributions. Instances of a single lesion affected ROI in one subject were excluded, as distribution-based comparisons were not possible. Where applicable p-values were computed using Fisher’s combined probability test.

## Results

The LeAPP pipeline and its integrated correction methods improved the resulting data quality as compared to the baseline HCP pipeline for brains with stroke lesion. This study further showed the ability of LeAPP for processing low quality structural and functional stroke patient data as no stroke patients had to be excluded due to lesion topology causing the processing pipeline to fail as previously reported on the same data set^6^, showing the direct relevance for clinical data. While global parcellation based volumetric agreement measures show no difference between processing pipelines (p-values: dice score=0.76, Jaccard score=0.49, volume difference=0.56; **Figure 4 (A)**), region-wise comparisons found significant improvements of LeAPP across all volumetric agreement measures for directly lesion affected and not-directly affected ROIs (p-values: dice score = 3.76e^-75^ and 0.0, volume difference = 4.36e^-19^ and 8.79e^-176^; **Figure 4 (C)**). We found a similar impact on the downstream processing step of creating structural connectomes. A significant difference in global average node strength was identified while the difference in clustering coefficient and average centrality do not show significance (p-values: node strength=3.09e^-7^, clustering coefficient=0.08, node centrality=0.19; **Figure 4 (B)**). Similar to the volumetric reconstruction quality measures for parcellations, we found a local impact of processing on network level as well with differences in node strength for directly lesion affected and not-directly affected ROIs and centrality for not-directly affected ROIs (p-values: node strength = 1.63e^-61^ and 0.0, node centrality = 0.25 and 0.02). **Table 1** presents all test statistics comparing reconstruction quality metrics for LeAPP and HCP structural processing of the validation ALE data set including effect sizes. While the lower number of directly lesion affected ROIs might contribute to the reduced effect sizes and lack of significance in e.g. ROI based node centrality compared to not-directly affected ROIs, an additional interesting finding of this study is that all local measures showed significance for the subset of not-directly lesion affected ROIs highlighting lesion impact in downstream processing beyond lesion location. The feasibility of the results is further strengthened when investigating differences in reconstruction metrics on individual subject level, as shown in **Figure 4 (E)**, displaying the largest differences between processing frameworks around the location of the lesion.

**Figure 4.**
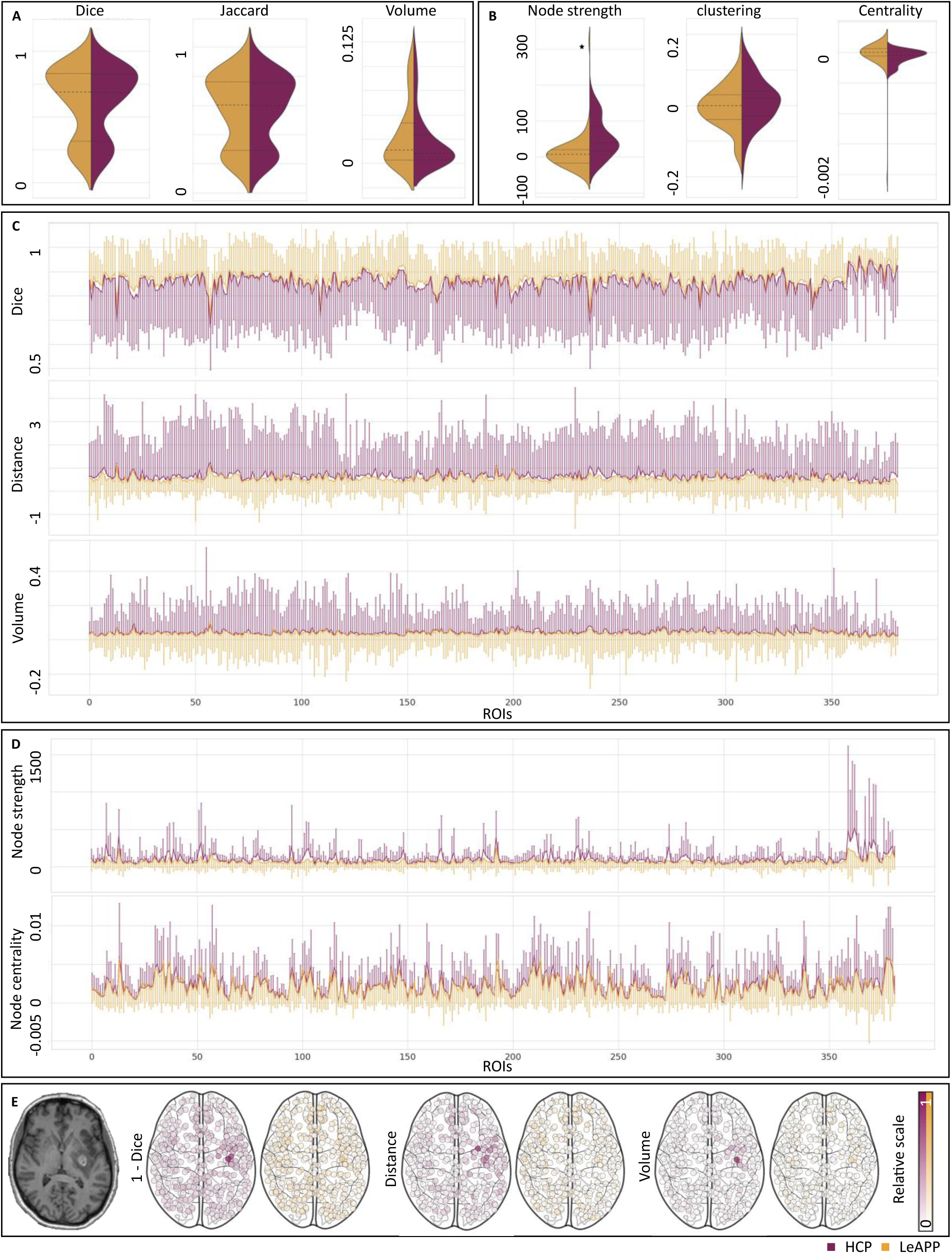
Reconstruction quality results. (**A**) Metrics for LeAPP (yellow) and HCP (purple) showing similar distributions across global measures. (**B**) Differences between LeAPP and HCP in global network metrics show a significant reduction in average global node strength difference between LeAPP and ground truth compared to HCP and ground truth differences. (**C**) Median local ROI based agreement metrics (line) with standard deviation shown as error bar. For visualization only one-sided error bars are displayed. (**D**) Median differences in network based local metrics for all ROIs also showing significantly smaller differences for LeAPP over HCP (see Table 1). Local differences highlight the increased sensitivity of ROI based measures in contrast to global measures based on full brain properties. (**E**) Exemplary ROI based agreement metrics for a single subject. Anatomical T1w image (left) shows the clear lesion artefact. The normalized agreement metrics reproduce the largest differences in processing around the focal lesion visible in the MRI image. Dice score is shown as 1 – dice score for consistency in visualizations.

**Table 1.**
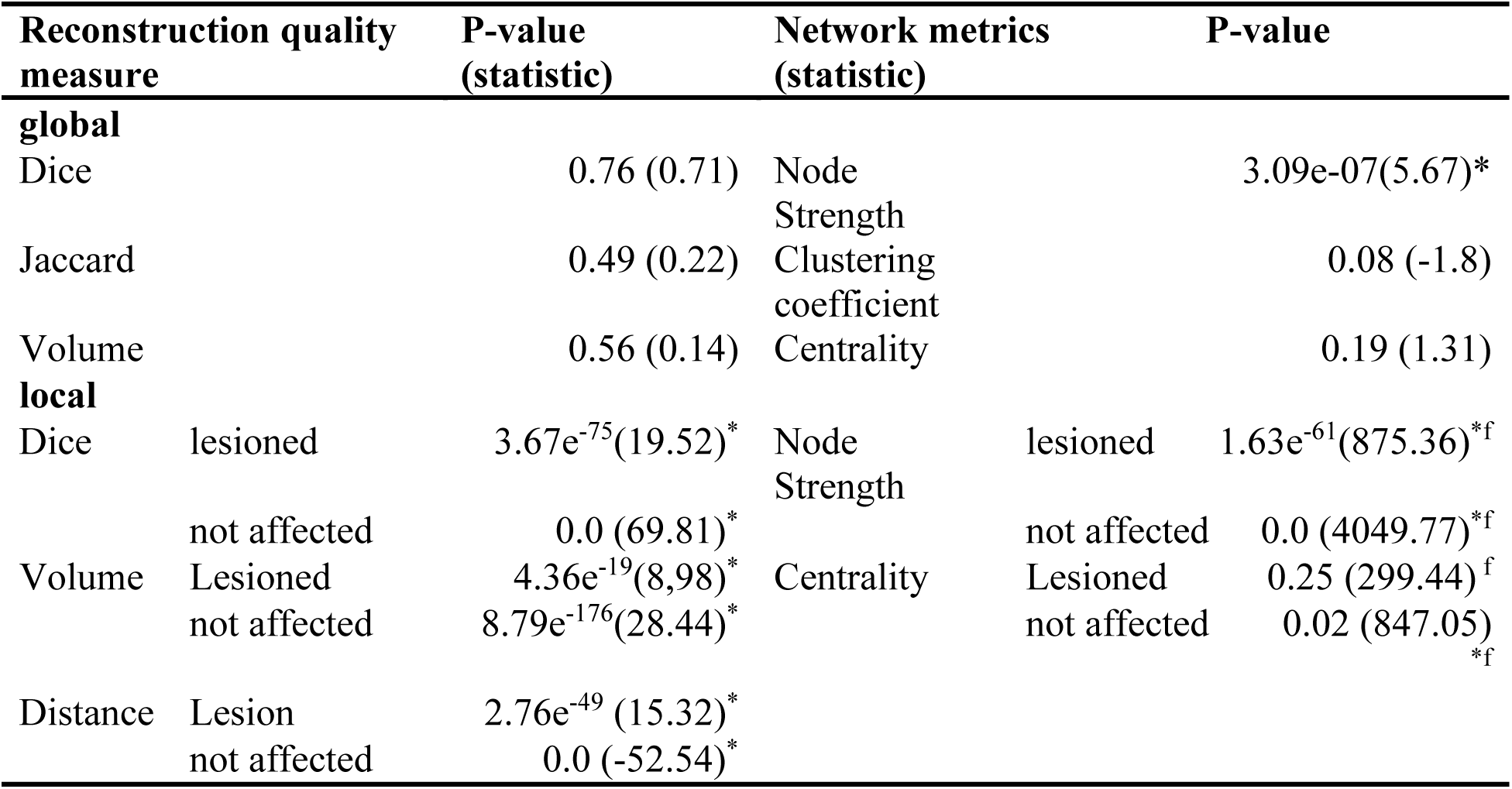
Test statistics for reconstruction quality. measures Differences between LeAPP and HCP processing were evaluated via reconstruction quality measures for created brain parcellations (left) and differences in network metrics for structural connectomes (right). Global measures did not show difference for brain parcellations but a significantly smaller difference in average global node strength between ground truth and LeAPP versus ground truth and HCP based connectomes. Local differences were significant for brain parcellations across measures. Differences in local network measures were significant for node strength for affected and not affected ROIS. Centrality did not show differences in lesion ROIs but overall, in not-affected ROIs. * Significance, f Fisher’s method

## Discussion

The objective of this study was to design and validate an automated reproducible processing pipeline that can specifically meet the challenges of processing stroke patient MRI data that also enables whole brain network simulation for stroke data using TVB.

In summary, we show that LeAPP leads to a significant improvement in recovering important structural brain information when processing MRI data that are affected by stroke lesions. The validation approach presented here shows that LeAPP is widely applicable from structural analysis to connectomics. An additional benefit of LeAPP is that it automatically creates TVB-ready data to facilitate accurate brain network modelling (BNM) for stroke. LeAPP further allows for introduction of ROI specific information such as lesion load, that had been previously linked to network disruption^31^, into individual BNMs. The following part of this paper moves on to discuss several aspects of such a multi-modal processing approach and presents remaining challenges for the field to enable accurate incorporation of lesion induced artefacts, while at the same time minimizing biases in processing and downstream analysis based on processing methods and data acquisition.

While MRI is the standard for noninvasively investigating brain tissue damage in research^2^, previous studies providing automated MRI processing frameworks^7,21,32,33^ are currently still limited to artifact free high resolution imaging data or specific processing steps^25,32^. The clinical context of data acquisition for acute stroke patients often severely limits the availability of scanning protocols. This leads to the common practice of manual adjustments and corrections during processing for each subject, reducing the automation potential, or applying pipelines initially used for different disease populations^2,6^. Our results show the advantage of LeAPP over such prior frameworks, as defined as necessary by^1^, in processing low quality functional imaging data (**Supplementary Figure S3**) as well as low resolution FLAIR imaging, which is often acquired instead of a T2 image in the clinical context of stroke^34^. It further represents, to the best of our knowledge, the first complete and automated multi-modal processing pipeline incorporating advanced correction methods, that have previously been stand-alone solutions^10,25,35^. These improvements, in addition to providing high quality reconstruction of volume and network information from MRIs, help to avoid excluding data due to poor quality or large artifacts, thereby facilitating research with larger cohort and longitudinal studies in the future.

### Limitations

Previous studies have shown an increase in accuracy of image registration^9,26^ when applying CFM to lesion data. Nevertheless, a residual distortion of processing results after applying CFM cannot be avoided^36^ leading to a baseline error in processing as found in the reduced agreement measures even in ROIs not directly affected by the lesion. Further research is needed to understand the impact of CFM and account for its effects on group level comparisons.

The extent of lesion abnormality in MRI images, such as flattened intensity value distributions (**Supplementary Figure S2**), has previously been shown to depend on various factors, such as imaging modality and relative time of acquisition to onset^37–39^. This creates the need for automated correction measures to enable appropriate processing for all modalities. While performing virtual brain transplant (VBT), as implemented in this study, offers such a correction for anatomical modalities like T1w, T2 and FLAIR, the application to EPI sequences, such as fMRI and DWI, is less feasible for recovering the information distorted in the affected region. A more thorough investigation of how such lesion abnormalities affect these modalities must be performed. In the current study we followed previous studies^40^ and excluded lesion signal from tractography by incorporating the corresponding lesion mask as pathological tissue in the context of anatomically constrained tractography. While we did not perform specific corrections to the fMRI lesion signal, the lesion induced abnormality did affect our decision not to perform independent component analysis (ICA) for denoising and motion correction (e.g., FIX-ICA;^41^). Previous studies have evaluated the robustness of such frameworks in the presence of stroke lesions^42^ showing a significant signal loss when applying automated complex approaches. It was further shown that lesion-based variance patterns differ significantly from healthy tissue, allowing for component-based lesion identification^43^ in resting-state fMRI. Further research into our understanding of signal and noise sources as identified by ICA at the focal lesion is needed in particular in the presence of task-based clinical fMRI data as present in the current study. The presented pipeline, LeAPP, currently remains limited to correction at the location of the lesion as defined by a segmentation mask that must be provided a priori. This limits its ability to correct for distortion and diaschisis effects caused by the damaged tissue and further research is needed to facilitate appropriate processing and analysis methods for incorporating distributed lesion impact. Furthermore, it requires time-consuming lesion segmentation by trained staff. A review by^44^ found that automated segmentation algorithms perform not yet sufficiently to justify integration into fully automated pipelines. While there have been several studies further developing such algorithms across diseases^45–47^ and species^48^, the performance for human stroke lesion segmentation did not yet improve to a level justifying the integration in LeAPP.

### Conclusions

In summary, we found that LeAPP represents significant progress for processing multi-modal neuroimaging patient data with lesion pathologies and its level of automation makes this workflow readily available to the scientific community. The pipeline can be used as a standard – well documented and versioned – tool for the processing of stroke imaging data thus ensuring a high degree of reproducibility and comparability of results. This is an important advancement as it can aid future studies in processing large cohorts of stroke data. Furthermore, LeAPP generates and outputs brain networks in TVB-ready formats that facilitate dynamic brain network modelling of virtual stroke brains with the TVB software. This will further facilitate simulation-based research into underlying recovery mechanisms, that so far have not been well understood.

## Acknowledgements

We thank Paul Triebkorn for fruitful initial discussions on the feasibility and implementation of the approach discussed in this study.

## Supplementary Material

Supplementary Materials is provided for visualizations and figures as well as methods and tables.

## Author Contributions

**PB:** Conceptualization, Data Curation, Formal Analysis, Investigation, Methodology, Software, Validation, Visualization, Writing - Original Draft Preparation, Writing - Review & Editing**, KD:** Methodology, Writing - Review & Editing**, AK:** Writing - Review & Editing**, MS:** Writing - Review & Editing**, JF:** Resources**, MB:** Resources**, RS:** Resources, Writing - Review & Editing**, BC:** Resources, **GT** Resources**, CG:** Resources, **PR**: Conceptualization, Funding acquisition, Supervision, Writing - Review & Editing, Management

## Nonstandard Abbreviations and Acronyms

ALE: artificial lesion embedding
BIDS: Brain Imaging Data Structure
CFM: cost function masking
DWI: diffusion weighted imaging
FLAIR: fluid attenuated inversion recovery
LeAPP: lesion aware automated processing pipeline
TVB: The Virtual Brain
VBT: virtual brain transplant

